# Mechanics regulate the postembryonic developmental maturation and function of sensory neurons in a pre-vertebrate chordate

**DOI:** 10.1101/2025.04.16.649091

**Authors:** Andreas Midlang, Carl Matthew Jones, Luc Seegers, Jorgen Hoyer, Sigurd Glørstad, Marios Chatzigeorgiou

**Author notes:** These authors contributed equally (Andreas Midlang, Carl Matthew Jones).

## Abstract

Physical forces are emerging as important contributors to a wide diversity of biological processes across multiple scales and contexts. The development of the nervous system is infuenced by mechanical forces exerted on neurons and glia by the surrounding environment. However, the extent to which tissue mechanics influence the function and morphology of the post-embryonic nervous system remains unclear. Here we leverage the post-embryonic larval period of the pre-vertebrate chordate *Ciona intestinalis* to study the dynamic interplay between nervous system mechanics, cellular and nuclear morphology and neuronal function.

During this post-embryonic period the larval head undergoes substantial stretching along the anterior-posterior axis. We show that this macroscopic morphological change is associated with a significant change in cell and nuclear shape (stretching) of the polymodal sensory neurons located in the papillar organs.

Using FRET based genetically encoded tension sensors we show that mechanical tension at the focal adhesions and the nuclear envelope of these papillar sensory neurons increases with time. At the functional level Ca^2+^ imaging experiments reveal that presentation and removal of a chemosensory stimulus that promotes settlement and metamorphosis elicits significantly stronger responses in late swimming larvae papillar neurons compared to those newly hatched ones. Finally, by combining optoGEF-RhoA dependent control of cellular forces and Ca^2+^ imaging we demonstrate that the strength of chemosensory responses and settlement behavior can be modulated by mechanical forces.

## INTRODUCTION

Our brains are amongst the softest tissues in the body(*3*), however, the nervous system is characterized by mechanical heterogeneity(*4, 5*). The mechanical milieu of the brain works together with biochemical signals to control key aspects of neuronal development such as cell migration, axon and neurite extension, cellular morphology, protein expression and repair response following injury(*6-10*).

The contributions of mechanics to the development of the nervous system are intensely studied increasing our understanding of the underlying mechanisms(*9-11*). However, it remains largely unclear whether cellular mechanics are important for neuronal physiology (e.g. sensory perception and information processing) and postembryonic changes in the nervous system.

Previous studies attempting to address these questions have found that the intrinsic mechanical properties of neurons and glia as well as those of surrounding tissues change with age and perturbation of the mechanical properties of brain tissue may lead to impaired sensory and cognitive ability(*12, 13*). In addition, micropatterned neuronal cell culture experiments revealed that stiff substrates promoted the formation of synaptic vesicles and enhanced electrophysiological activity of neural networks(*14*). Finally, the most convincing evidence for the impact of cellular mechanics on neuronal function comes from an *in vivo* study using *C. elegans* which demonstrated that the cytoskeletal protein Spectrin is held under constitutive tension in touch neurons. Loss of Spectrin-dependent tension selectively impaired touch sensation, advocating that pre-tension is required for the ability of touch neurons to respond effectively to external mechanical cues(*15*).

Despite these studies, direct observation and manipulation of neuronal activity and mechanical forces *in vivo* remains a formidable challenge and how exactly changes in nervous system mechanics during post-embryonic development influence neuronal activity remains an important gap in our knowledge(*16*).

The pre-vertebrate chordate *Ciona intestinalis* offers a potentially interesting paradigm to address this knowledge gap. During the first ∼6 hours (at 18°C) of its post-embryonic life the freely swimming larva undergoes a striking morphological transformation whereby its trunk shape changes from spherical to a spindle-like shape(*1*). This change is thought to coincide with significant changes in the internal tissues of the larva including the nervous system, which prime the animal for the next stages of its life-cycle, namely the processes of larval settlement and metamorphosis(*1*).

Here we show that the change in trunk geometry is associated with morphological changes in cell and nuclear shape of the sensory neurons located at the adhesive papillae. By using the genetically encoded tension sensors NesprinTS and VinculinTS we find that mechanical tension increases at the nuclear envelope and focal adhesions respectively of the papillae as the larvae progress from the hatching to late swimming stage. These changes in mechanical tension are associated with a change of Ca^2+^ responses to chemosensory stimuli. Finally, we employ optogenetic tools capable of altering cellular contractility to demonstrate that an increase in mechanical tension enhances chemosensory responses while the opposite outcome is achieved when we decrease mechanical tension.

## RESULTS

The post-embryonic period from hatching larva to late swimming larva was recently divided into four anatomically distinguishable stages that reflect at least in part the changes observed in trunk shape(*1*). However, quantitative descriptions of larval trunk shape and nervous system morphology changes over time are missing.

Therefore, we reared transgenic *Ciona* embryos at 18°C and fixed larvae at four different post-embryonic stages (St. 26 to St. 29) to quantify trunk morphometrics and used a reporter of peptidergic neurons to visualize any changes in the structure of the nervous system (Fig. 1A, B and C).

**Fig. 1.**
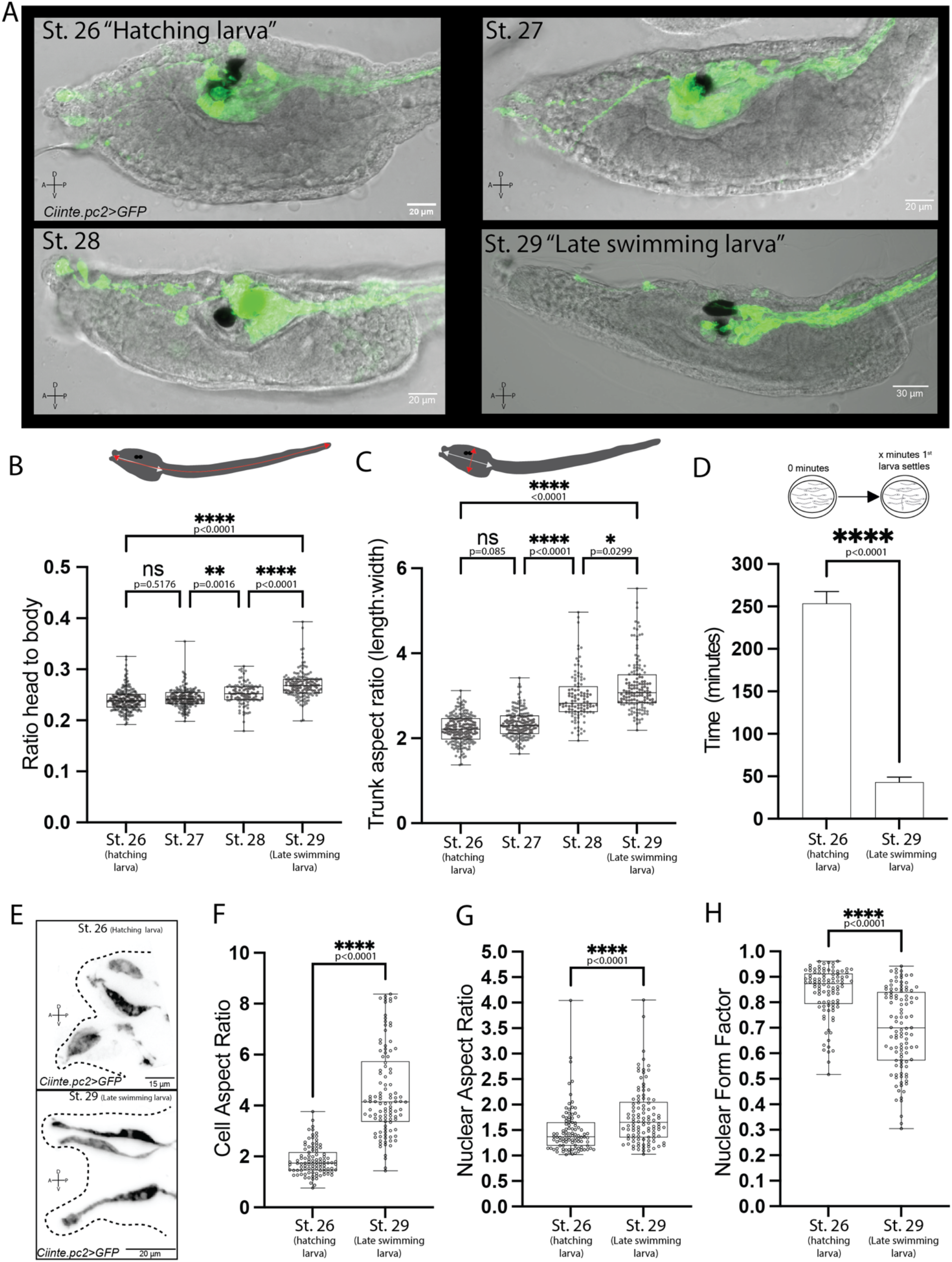
Post-embryonic larval development leads to dramatic changes in trunk shape and sensory neuron cellular and nuclear morphology. (A) Confocal micrographs of Ciona larvae at the four different developmental stages as defined by Hotta et al. St. 26 corresponds to hatching larva, while St. 29 refers to late swimming larva(*1*). The peptidergic neurons of the nervous system are labelled by *Ciinte*.*pc2*>GFP. Scale bars=20μm (St. 26-St. 28 or 30μm for St. 29). (B) Quantification of head to body ratio change during post-embryonic larval development. 200≥n≥103 larvae per stage. (C) Quantification of trunk aspect ratio across larval post-embryonic development. 197≥n≥106 larvae per stage. For panels C and D we performed a Kruskal-Wallis test followed by Dunn’s multiple comparisons test. (D) Quantification of settlement onset for freshly hatched larvae and late swimming larvae. 22 assay plates per stage, n≥50 per assay. For statistical analysis we performed a Mann-Whitney test. (E) Confocal micrographs of the main sensory cell type in the Ciona papillae termed Papillae Sensory Neurons (PSNs), labelled by *Ciinte*.*pc2>GFP*. Top and bottom panel shows the PSNs of a hatching larva, while the bottom panel (F-H) Box plots quantifying the cell aspect ratio (E), nuclear aspect ratio (F) and nuclear form factor (G) of Papillae Sensory Neurons (PSNs) in hatching (St. 26) and late swimming larvae (St. 29). n=98 (St. 26) and n=106 (St. 29). For statistical analysis we performed a Mann-Whitney test.

We found that the head to body ratio (Fig. 1B) and trunk aspect ratios (Fig. 1C) were significantly higher as *Ciona* larvae progressed through their postembryonic development confirming previous qualitative observations(*1*).

Previous studies have suggested that the observed trunk shape changes are associated with the acquisition of the ability to settle and metamorphose (the term used was “metamorphic competence”)(*17, 18*). We tested this hypothesis by measuring the time it takes for freshly hatched or late swimming larvae to settle after being transferred to a plastic Petri dish. We found that the median time it took for hatching larvae to start the settlement process was 262.5 minutes, in contrast to late swimming larvae which took 30.0 minutes (Fig. 1D). We repeated the experiments in the presence of 10mM NH_4_Cl which is a cue that can accelerate settlement and metamorphosis(*19*). We discovered that hatching larvae populations require a median time of 270 minutes for the first larva to settle (fig. S1A). In contrast, in populations of late swimming larvae the presence of 10mM NH_4_Cl accelerated the onset of settlement from a median of 30.0 minutes to a median of 15.0 minutes (fig. S1A). Our results suggest that only late swimming larvae are primed to settle in the presence of mechanical and chemical signals.

Previous studies had made qualitative morphological observations indicating that the papillae of the larvae transition from a pyramidal/oval-like shape (immature papillae) to an elongated shape (prior to adhesion)(*1, 20*). We quantified the aspect ratio of Papillae Sensory Neurons (PSNs, labelled with *Ciinte*.*pc2>GFP*) in hatching larvae (St. 26) and late swimming larvae (St. 29). We found that late swimming larvae PSNs were characterized by a significantly higher aspect ratio compared to hatching larvae (Fig. 1E). In addition, we quantified the length of the sensory endings (distal endings) of PSNs and axon lengths between PSNs and the upstream RTENs (fig. S1B-S1E). We found that both the sensory endings (fig. S1D and S1E green arrowheads) and axon lengths were significantly larger in late swimming larvae (fig. S1B and S1C). It is known that the nucleus can be deformed in response to mechanical force and shape changes that occur at the cell periphery(*21, 22*). We thus wondered whether the shape changes of the PSNs were associated with changes in the nuclear shape. We found that nuclei of late swimming larvae PSNs had a higher aspect ratio and significantly lower nuclear form factor (otherwise known as nuclear circularity or contour ratio(*23, 24*)) compared to hatching larvae PSNs (Fig. 1F and G). This suggests that as the larvae transition through their postembryonic development the nuclei elongate, like the PSNs cells and adopt less round contours that have lobular irregularities.

A substantial body of studies has shown that cell and nuclear shape changes are associated with changes in mechanical forces. To quantify the mechanical forces acting at focal adhesions and the nucleus we leveraged the FRET-based genetically encoded tension sensors VinculinTS(*25*) and NesprinTS(*26*) respectively (Fig. 2A and B). We examined FRET of the VinculinTS and NesprinTS tension sensors in hatching larvae papillae as compared to late swimming larvae papillae (Fig. 2C). We found that the papillae of late swimming larvae labelled with VinculinTS or NesprinTS exhibited significantly higher FRET index values compared to those of hatching larvae (Fig. 2D and E). This suggests that as the post-embryonic developmental progression is associated with an increase in tension at focal adhesions and the nuclear envelope. The papillae cells of the late larvae were characterized by a significantly higher concentration of actin filaments at their apical tips compared to those of early larvae (Fig. 2F-I; fig. S2A-S2D).

**Fig. 2.**
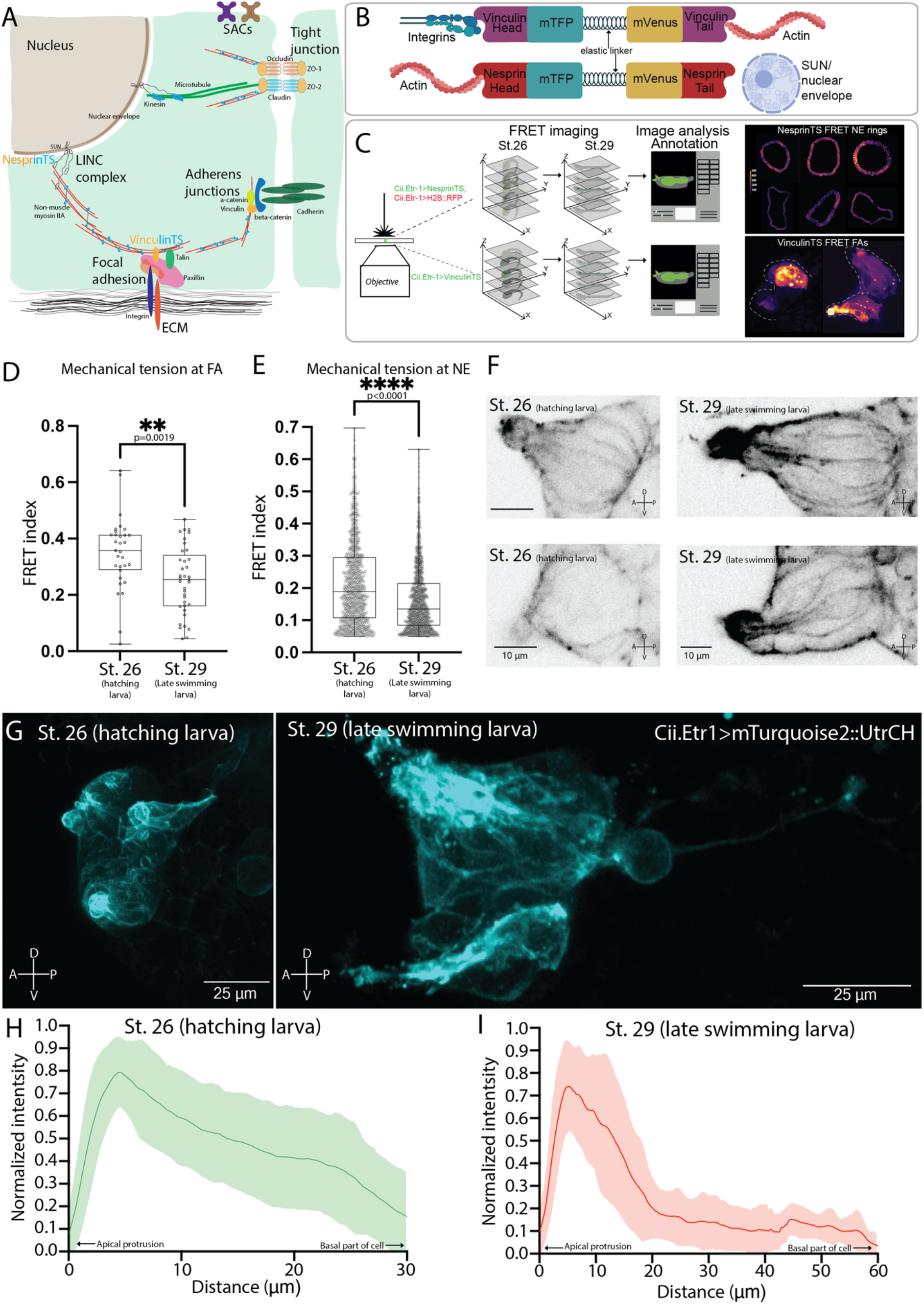
Post-embryonic development of papillae sensory cells experience an increase in mechanical tension and a reorganization of the cytoskeleton. (A) Schematic of a cell highlighting some of the key sites and players of mechanotransduction/mechanosensation. The site of localization of the VinculinTS and NesprinTS sensors are highlighted. (B) Schematic of molecular organisation of the two tension sensors used in our study. (C) Pipeline for FRET measurements of mechanical tension using the NesprinTS and VinculinTS sensors. Confocal stacks were recording at St.26 and St.29 larval stages. To identify neuronal nuclei we used the marker *Ciinte*.*Etr-1>H2B::RFP*, which were segmented with Cell Pose 2.0(*2*). (D) Quantification of FRET index recorded from VinculinTS at St.26 (n=31 animals) and St.29 (n=37 animals). (E) Box plots quantifying the average NesprinTS FRET index around the nuclear membrane of papillae sensory cell nuclei of St.26 (n=1322 nuclei) and St.29 (n=1365 nuclei). For statistical analysis of data shown in panels C and D we performed Mann-Whitney tests. (F) Confocal micrographs of papillae from St.26 and St.29 stained with an antibody against F-actin. Scale bars=10μm. (G) Representative maximal projection confocal micrographs of papillae expressing *Ciinte*.*Etr-1>mTurquoise::UtrCH* during St.26 (left micrograph) and St.29 (right micrograph). Scale bars= 25μm. (H, I) Average traces of normalized mTurquoise::UtrCH intensity across papillae sensory cells (Apical protrusion-left end of graphs, Basal part of cell-right end of graphs) for St.26 (panel H n=40; 10 larvae) and St.29 (panel I, n=56 cells, 14 animals). The solid line in each trace correspond to the mean normalized intensity and the shaded area corresponds to the Standard Deviation.

We wondered whether a change in the shape and mechanical tension of the papillae results in a change in the Ca^2+^ responses of the PSNs in response to sensory stimulation. We have previously shown that the PSNs (and the neighbouring axial columnar cells, ACCs) respond to various sensory cues include chemosensory stimuli such as NH_4_Cl(*19*). We expressed the genetically encoded Calcium indicator GCaMP6s under the *Ciinte*.*pc2* promoter and imaged Ca^2+^ responses to 10mM NH4Cl from the PSNs of hatching larvae (St. 26) and late swimming larvae (St. 29) (Fig. 3A and B; Movies S1 and S2). Activation of the PSNs with 10mM NH4Cl elicits strong stimulus on and stimulus off responses in late swimming larvae (Fig. 3C solid red line; fig. S3B), in stark contrast to hatching larvae (Fig. 3C solid green line; fig S3A) which show a significantly smaller stimulus on response (as quantified by peak amplitude and area under the curve) and they lack a pronounced stimulus off response (Fig3C and D).

**Figure 3.**
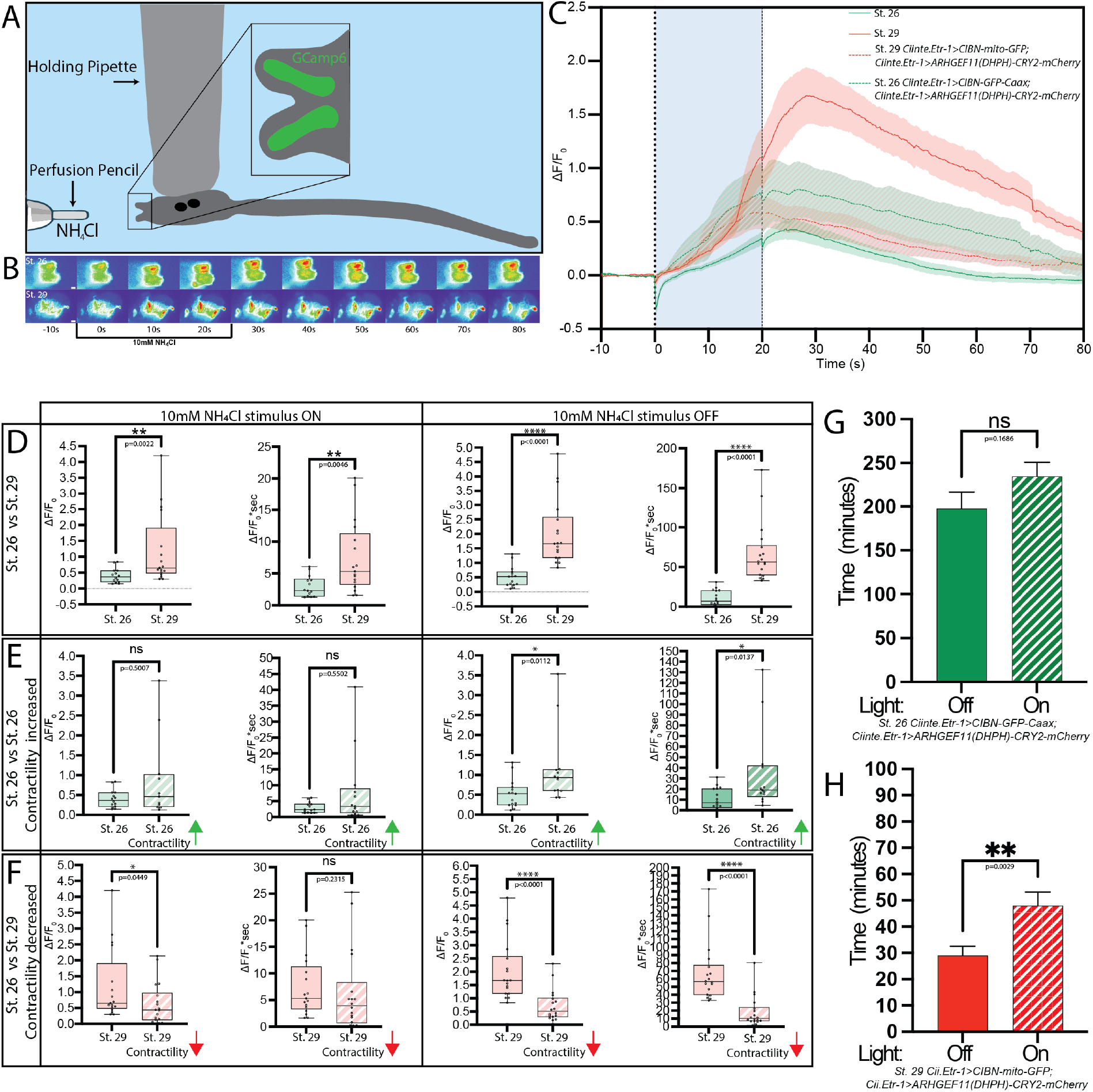
Chemosensory responses of PSNs increase with postembryonic maturation and are sensitive to optogenetically induced changes in cell stiffness. (A) Schematic of the apparatus used to immobilize and stimulate *Ciona* larvae with 10mM NH_4_Cl while recording calcium activity from PSNs. (B) Montage of two example movies from St.26 and St.29 larvae, showing PSN cells expressing GCaMP6s under the *pc2* promoter responding to 10mM NH_4_Cl. (C) Average traces of PSN calcium responses to 10mM NH_4_Cl recorded from St.26 larvae (solid green trace), St.29 larvae (solid red trace), St.26 larvae expressing the optogenetic tool components *Ciinte*.*Etr-1>CIBN-GFP-Caax;Ciinte*.*Etr-1>ARHGEF11(DHPH)-CRY2-mCherry* which cause an increase in cellular tension upon illumination (dashed green trace) and St.29 larvae expressing the optogenetic tool components *Ciinte*.*Etr-1>CIBN-mito-GFP;Ciinte*.*Etr-1>ARHGEF11(DHPH)-CRY2-mCherry* which cause an decrease in cellular tension upon illumination (dashed red trace). The solid lines indicate mean values and the shaded areas indicate Standard Error of the Mean. (D-F) Quantification and comparison of calcium responses peak features to 10mM NH_4_Cl stimulus presentation (ON) and removal (OFF) in St.26 and St.29 animals (D), St.26 and St.26 expressing and activating *Ciinte*.*Etr-1>CIBN-GFP-Caax;Ciinte*.*Etr-1>ARHGEF11(DHPH)-CRY2-mCherry* (E), and St.29 against St.29 expressing and activating *Ciinte*.*Etr-1>CIBN-mito-GFP;Ciinte*.*Etr-1>ARHGEF11(DHPH)-CRY2-mCherry*. (G) Quantification of settlement onset for St.26 larvae and St.26 *Ciinte*.*Etr-1>CIBN-GFP-Caax;Ciinte*.*Etr-1>ARHGEF11(DHPH)-CRY2-mCherry*. 30 assay plates per condition, n≥50 per assay. (H) Quantification of settlement onset for St.29 larvae and St.29 *Ciinte*.*Etr-1>CIBN-mito-GFP;Ciinte*.*Etr-1>ARHGEF11(DHPH)-CRY2-mCherry*. 30 assay plates per condition, n≥50 per assay. For statistical analysis we performed a Mann-Whitney test. For statistical analysis of data presented in panels D-H we carried out Mann-Whitney tests.

We then asked whether a change in mechanical tension in the papillae would alter the Ca2+ responses observed in hatching and late swimming larvae. To modulate mechanical tension in the papillae we used an optogenetic tool that relies on controlling the subcellular activation of RhoA using a CRY2/CIBN light-gate dimerization approach. Translocation optoGEF-RhoA to the plasma membrane causes an increase in contractility. By contrast, light-induced translocation of optoGEF-RhoA to mitochondria results in reduction in contractility. We generated two types of transgenic larvae expressing *Ciinte*.*pc2>GCaMP6s* either with the plasmid cocktail that would increase contractility in hatching larvae *(Ciinte*.*Etr-1>CIBN-GFP-Caax;Ciinte*.*Etr-1>ARHGEF11(DHPH)-CRY2-mCherry)* or with the plasmid combination that would decrease contractility in late swimming larvae (*Ciinte*.*Etr-1>CIBN-mito-GFP;Ciinte*.*Etr-1>ARHGEF11(DHPH)-CRY2-mCherry)*. Optogenetic activation was provided by the light-source that was also used to excite GCaMP6s. When stimulated with 10mM NH_4_Cl the PSNs of hatching larvae elicited stimulus on responses that appeared larger than those of hatching larvae without the optogenetic tool (Fig. 3C green dashed line; fig. S3C, S3E; Movie S3). Quantification of peak features such as peak amplitude and area under the curve showed that contractility increases because an optogenetically induced increase in cellular contractility enhanced significantly the stimulus off response but not the stimulus on response (Fig. 3E).

Stimulation of late swimming larvae carrying the optoGEF-RhoA tool mix that leads to contractility decrease showed a significant reduction in both stimulus on and stimulus off responses (Fig. 3C, 3D; fig. S3D, S3F; Movie S4). During the stimulus on period contractility decrease led to a significant reduction in Ca^2+^ peak amplitude (but not area under the curve), while when the NH_4_Cl stimulus was removed we observed a very large reduction in both peak amplitude and area under the curve (Fig. 3C and F). To determine if optogenetic manipulation of tension in the papillae influenced behavior, we carried out a settlement assay in the presence of 10mM NH_4_Cl using either hatching larvae expressing *Ciinte*.*Etr-1>CIBN-GFP-Caax; Ciinte*.*Etr-1>ARHGEF11(DHPH)-CRY2-mCherry* or late larvae expressing *Ciinte*.*Etr-1>CIBN-mito-GFP;Ciinte*.*Etr-1>ARHGEF11(DHPH)-CRY2-mCherry*. We found that increase of contractility did not significantly promote settlement onset (Fig. 3G). In contrast, a decrease in contractility in late larvae significantly increased the time taken to initiate settlement (Fig. 3H).

Taken together our data show that the chemosensory neurons of hatching larvae and late swimming larvae show significantly different Ca^2+^ response profiles in response to a sensory cue. In addition, modulation of cellular contractility and mechanical tension can alter Ca^2+^ responses and behavioral output.

## DISCUSSION

Here we have demonstrated that during postembryonic development, sensory neurons in *Ciona intestinalis* larvae experience changes in morphology, mechanical tension levels and ultimately in their chemosensory responses. We have offered evidence that these chemosensory responses can be modulated by a change in cellular contractility.

Measurement of larval trunk, body length and width allowed us to observe significant changes in trunk and body shape during postembryonic development. We found that this morphological transition underlies the ability of the larvae to settle and metamorphose. It is therefore conceivable, that reaching a stretch threshold along the body length axis acts a primer for settlement and metamorphosis. Interestingly, a recent study using *C. elegans* proposed a similar mechanism which triggers the L1-L4 larval-stage transitions in the nematode(*27*).

We have found that the morphological changes in trunk shape are accompanied by changes in cellular and nuclear morphology of the sensory cells found in the larval adhesive organs (the papillae). Vertebrate neurons in the peripheral and central nervous system are not only functionally but also morphologically diverse at the level of cellular and nuclear morphology and architecture(*28-30*). Changes in the cellular and nuclear morphology of neurons have mostly been associated with disease or aging(*29, 31-34*). In *Ciona* larvae, changes in neuronal cellular and neuronal morphology are unlikely to be directly related to disease or aging due to the limited span over which these modifications take place (i.e. 6 hours at 18°C). Alternative mechanisms that could induce these morphological modifications include neuronal activity and mechanical forces. For example, it has been shown that the nuclei of cultured hippocampal neurons infold in an activity-dependent manner(*35*) and there is an increasing body of evidence (especially ex-vivo) that demonstrate an important role for mechanical forces in shaping neuronal cellular and nuclear morphology under physiological or pathological conditions(*6, 9-11, 36*).

Our experimental observations show that these morphological alterations are associated with a significant increase in mechanical tension at both focal adhesions and the nuclear envelope of papillae sensory cells as they “mature” during post-embryonic development. To date this phenomenon has been described in a handful of studies utilizing mammalian species. One study reported that postnatal brain maturation in mice is associated with a shift of brain mechanical properties towards a more rigid behavior(*37, 38*), while another report following the postnatal development of the ferret brain ascertained an increase in overall brain stiffness with time(*39*). Taken together our work suggests that postembryonic increase in brain stiffness may be a more widespread mechanism that contributes to chordate brain maturation than previously believed.

To date the cross-modal interplay between sensory stimulus-driven neuronal activity and mechanical signals has only been convincingly demonstrated for vision. In *Drosophila* photoreceptors, where absorption of light leads to the activation of a signalling cascade which changes the physical properties of the cell membranes. Therefore, microvilli respond to light by contracting. The perceived increase in mechanical forces opens “light-sensitive” channels, which are, in fact, mechanosensitive(*40*). Likewise, a study in Xenopus rod cells demonstrated that the mechanosensitive channels TRPC1 and Piezo1 are crucial for photoreceptor function, suggesting that mechanosensitivity is a critical component of vertebrate vision(*41*). Here we have shown that a second sensory modality, namely chemosensation, is modulated by mechanical forces. The precise mechanism by which this happens remains unclear, but several hypotheses can be raised. Our optogenetic experiments show that acute manipulation of cellular stiffness can alter chemosensory responses. Thus, *Ciona* may sense chemosensory cues through polymodal ion channels (e.g. TRPs) which can respond to both chemical and mechanical cues(*42*). A change in cell stiffness could alter the conformation of these channels, and thus, enhance or dampen their response to a chemical cue. An alternative yet complementary mechanism would involve mechanically induced deformations of the nucleus (which we do observe) leading to changes in gene expression of molecules involved in sensory transduction and signal amplification. In line with this hypothesis, a previous study reported that “competent” larvae stained more strongly for the β-Adrenergic receptor compared to hatching larvae and that noradrenergic signalling promoted settlement and metamorphosis(*43*). In addition, we found that a contractility increase in hatching larvae had a limited impact on chemosensory responses (in contrast to contractility decrease in late larvae) suggesting that possibly key molecular components of the transduction machinery were either not expressed or not functional at that early stage of postembryonic development.

Finally, we demonstrated that a decrease in mechanical tension in the papillae of late swimming larvae delays the onset of settlement behavior, even if the settlement promoting chemical NH_4_Cl is present(*19*). Studies across multiple marine invertebrate taxa including sponges, corals, annelids, molluscs and ascidians pointed to the importance of gene expression changes associated with larval competence (*44-48*). Our study demonstrates that also mechanical forces are critical regulators of larval competence adding a new layer of complexity to this process of extreme importance for the ecology of our oceans.

## Acknowledgments

We would like to thank members of the Chatzigeorgiou lab for valuable feedback on the manuscript.

## Funding

Research Council of Norway 339399 (MC)

Research Council of Norway 234817 (To the Michael Sars Centre)

## Author contributions

Conceptualization: MC Methodology: AM, CMJ, MC

Investigation: AM, CMJ, MC, LS, SG

Visualization: AM, CMJ, MC, LS, SG

Funding acquisition: MC

Project administration: MC

Supervision: MC

Writing – original draft: MC, AM, CMJ

Writing – review & editing: MC, AM, CMJ, LS, SG

