## Supplemental Materials for "Mechanics regulate the postembryonic developmental maturation and function of sensory neurons in a pre-vertebrate chordate"

**Andreas Midlang et al.**

This file includes:

Materials and Methods

Figs. S1 to S3

Other Supplementary Material for this manuscript includes the following:

Movies S1 – S4

### Materials and Methods

#### Animal collection and rearing conditions

We collected gravid adult *C. intestinalis* from Døsjevika, Bildøy Marina AS, postcode 5353, Bergen, Norway. The relevant GPS coordinates are the following: 60.344330, 5.110812. We kept animals in a purpose-built facility with running sea water which had a temperature of 10°C and a pH of 8.2. The animals were kept under constant illumination to prevent spawning.

#### Egg fertilization, embryo, and larval rearing conditions

Egg collection, fertilization, and embryonic development conditions were carried out as previously described(*49, 50*) with the exception of the rearing temperature that was set to 18°C. Briefly we sacrificed two to three gravid Ciona adults per fertilizations. Eggs and sperm were collected seperately. 500 to 1000 eggs were spread out in a 9cm agarose coated petri dish containing ASW. 100μl of Tris activated sperm was added to the eggs and the fertilization was allowed to proceed for 15 minutes. Once fertilized, eggs were transferred to fresh 9cm plates and incubated in 18°C throughout the experiments.

#### Fixation, phalloidin staining and slide mounting

Larvae were added to the fixation buffer (MEM-FA (100mM MOPS pH 7.4, 500 mM NaCl, 2 mM $MgSO_4$, 0.5% Triton X-100 (v/v), 3.7% formaldehyde methanol free) and fixed for 15 min in RT. Then they were quenched in (1x PBS, 50 mM $NH_4Cl$, 0.35% Triton x-100) for 20 min at RT. We washed three times with 1xPBS for 5 minutes each. For phalloidin staining we incubated larvae in Alexa Fluor 647 Phalloidin for 2 days in PBS. Lastly, PBS was removed, and 3 droplets of mounting solution were added. Animals were then mounted on a slide with double-sided tape and covered with a coverslip. Regarding the tail-trunk measurements, animals were only fixed with PBST 0,1% once, and two times with PBS. Tail and trunk related measurements were performed using micrographs collected using a Zeiss Axiocam 506 color and the 10x/0.30 objective. These snapshots were then analyzed using Fiji(*51*).

#### Electroporation of Ciona zygotes

Electroporations were performed as previously described(*19*), with the following modification: We electroporated up fo 10μg per optogenetic construct, while for all other constructs we electroporated at 50 to 150μg of each construct. In all cases we used a capacitance of 1200μF.

#### Molecular cloning

To generate the tension sensors we subcloned VinculinTS(*25*) (a gift from Martin Schwartz (Addgene plasmid # 26019 ; http://n2t.net/addgene:26019 ; RRID:Addgene_26019) and NesprinTS(*52*) (a gift from Daniel Conway (Addgene plasmid # 68127 ; http://n2t.net/addgene:68127 ;

RRID:Addgene_68127) to the Gateway middle position vector using BP Clonase II. To generate the reporter for cytoskeleton distribution we subcloned mTurquoise2-UtrCH(*53*) (a gift from Dorus Gadella, Addgene plasmid # 98824 ; http://n2t.net/addgene:98824 ; RRID:Addgene_98824) to the 2nd position of Gateway using BP clonase II. The optogentic tools were constructed by subcloning the following: ARHGEF11(DHPH)-CRY2-mCherry (a gift from Xavier Trepat Guixer (Addgene plasmid # 89481 ; http://n2t.net/addgene:89481 ; RRID:Addgene_89481))(*54*), mito-CIBN-GFP (a gift from Xavier Trepat Guixer (Addgene plasmid # 89480 ; http://n2t.net/addgene:89480 ; RRID:Addgene_89480))(*54*) and pCIBN(deltaNLS)-pmGFP (a gift from Chandra Tucker (Addgene plasmid # 26867 ; http://n2t.net/addgene:26867 ; RRID:Addgene_26867))(*55*). These were then recombined with a *Ciinte.Etr-1* promoter in 1st position(*56*) and a unc-54 3'UTR(*56*) in the 3rd position with the pDESTII backbone using LR Clonse II. Expression vectors were purified using a Machery Nagel MidiPrep kit.

**Cell and nuclear morphology analysis.**

Animals electroporate with *Ciinte.Etr-1>Lck::mScarlet* (70μg) and *Ciinte.Etr-1> 3xnls::mNeonGreen* (50μg) were allowed to grow until hatching larva stage or late swimming stage at which points they were fixed for 15 min at room temperature (RT) in MEM-FA buffer (100 mM MOPS pH 7.4, 500 mM NaCl, 2 mM MgSO_4, 0.5% Triton X-100, 3.7% formaldehyde). Samples were quenched (PBS, 50 mM NH_4Cl, 0.35% Triton X-100) for 20 min at RT and washed three times in PBS. Animals were then mounted onto slides with mounting solution and double-sided tape, covered with a coverslip. Animals were imaged at an OLYMPUS FV3000 point-scanning confocal using a 40x silicon oil objective, with scanner set to 1024x1024 pixels. For illumination we used the following lasers 488 nm (max output 20 mW), 561 nm (max output 20 mW). For detection we used GaAsP photomultipliers. Data was loaded in ImageJ and shape outlines were drawn. For cells we quantified the aspect ratio, while for nuclei we calculated the aspect ratio and the form factor: $(4*\pi*\text{Area})/\text{Perimeter}^2$.

**VinculinTS FRET-based tension measurements**

Transgenic animals harbouring *Ciinte.Etr-1>VinculinTS* (electroporated at 100 mg) *at* either St.26 or St.29, were mounted on Mattek 35mm dishes with No.0 coverslips and immobilized using 0.03% MS-222. We imaged our animals on a FV3000 LSM microscope using a 40x silicon oil objective. We acquired 8 planes per animal at 800 × 800 pixels resolution. We measure the intensity of two channels, mTFP and mVenus excited by a 445 nm laser, along with a transmitted light channel. To account for signal bleed-through from mTFP to the mVenus channel, we measured the bleed-through factor by using a mTFP monomer. If we give the actual emitted intensity of the mTFP channel as $I_{mTFP}$, and the observed signal strength from the microscope for the channel as $C_{\mathrm{mTFP}}$, and similarly for mVenus, we get that,

$I_{mVenus} = C_{mVenus} - BC_{mTFP}$, where $B$ is the bleed-through factor that depends on the microscope, while $I_{mTFP} = C_{mTFP}$. To account for the level of expression of the sensor, we give the normalised FRET intensity, $F$, as $F = \frac{C_{mVenus} - BC_{mTFP}}{C_{mTFP}}$, therefore, has no units. To acquire regional FRET values the images were annotated using custom made software developed by the lab. This gives the user the ability to stage the animals into different Hotta stages and to mark the different regions of the nervous system (i.e. in this case we labelled the papillae). The regions (i.e. papillae) and stages are saved as data to each frame of the image to extract FRET values.

**NesprinTS FRET-based tension measurements**

Transgenic animals harboring *Ciinte.Etr-1>NesprinTS; Ciinte.Etr-1>H2B::RFP* transgenes (electroporated at 100µg and 40µg respectively) at either St.26 or St.29, were mounted on Mattek 35mm dishes with No.0 coverslips and immobilized using 0.03% MS-222. We imaged mVenus and mTFP channel along with H2B-RFP to detect nuclei. This was done on a SpinSORA microscope using a 40x silicon oil objective. We take 2,048 x 2,048 pixel images with stack thickness of 10 planes. As the Nesprin results are only meaningful around the nuclei we must first isolated these regions. We trained a CellPose(*2*) model to detect nuclei in the 3D-image and for each of the detected nuclei we proceeded with the central plane. We performed a binary dilation with a circle element with radius 4 to create a larger masked region around the nuclei, and then an exclusive-or of the masks to select only the pixels in the area around the nuclei. We then calculated the mean FRET intensity in each of the rings. The regions (i.e. papillae) and stages were defined using the same custom software that we used for the VinculinTS experiments.

**Utrophin localization experiments**

*Ciinte.Etr-1>*mTurquoise2::UtrCH was electroporated at 100µg and *Ciinte.Etr-1>Lck::mScarlet* at 60µg. St. 26 or St. 29 larvae were mounted on Mattek 35mm dishes with No.0 coverslips and immobilized using 0.03% MS-222. Animals were imaged using an OLYMPUS FV3000 LSM. We excited mTurquoise2 using the 445nm laser and mScarlet using 561nm. We acquired stacks at 1024 x 1024 pixels using a 40x Silicon Oil objective. The same laser intensity, detector gain and no offset were used for all animals imaged. We then generated sum projections in Fiji(*51*) and used the line tool (thickness 22) to measure the intensity profile from the apical to the basal side of each cell in the pappilae. Each intensity profile was normalized to account for expression variability.

**Calcium imaging**

St. 26 or St. 29 larvae were transferred in 99.97% ASW, with pH 8.4 at 14°C and 32% salinity, and 0.03% MS-222 for immobilization, in a 9 cm petri dish. To immobilize the larvae under the Axioskop A1 microscope, first we identified them using a 5x air objective. We then applied gentle suction using a customized glass pipette, which had an outer diameter of 120 µm and an inner diameter of 25 µm. The pipette was able to hold onto ~50% of the larval trunk (typically the ventral or lateral side) keeping the larvae in position for the chemical stimulation. We illuminated the larvae using a mercury lamp. The following combination of filters were used: BP470/20, FT493, BP505-530. We then switched to a 40x water immersion objective mounted on the Axioskop A1 microscope which was equipped with a Hamamatsu Orca Flash 4.0V2 CMOS camera. The camera field of view was ~320µm in each direction and the resolution was 2048x2048 pixels. We acquired the fluoresence data at 40Hz using the Hamamatsu Corporation imaging software tool HCImageLive. To achieve these frame rates and reduce file size we cropped the FOV using a bounding box around the papillae cells in HCl. The perfusion pencil that we used to deliver ASW and 10mM $NH_4Cl$ stimuli to the larvae was placed ~250 µm away from the papillae with the outlet pointing directly towards this structure. Larvae were initially exposed to ASW flow for 10 seconds. This first period was followed by 20 seconds of stimulation with 10mM $NH_4Cl$, followed with a switch back to ASW for another 50 seconds. For the optogenetic stimulation performed in the experiments shown in Figure 3, we used the same fluorescence light source as for exciting GCaMP6s. Continuous illumination Calcium imaging analysis was performed as previously described(*19*) using the software Mesmerize(*56*). For statistical analysis max peak amplitude and area under the peak data were plotted and analyzed using GraphPad Prism 10. First we performed a normality check using the Shapiro-Wilk test. This was followed performed a Mann-Whitney test. The number of values used in the analysis corresponds to individual traces.

**Settlement onset assays without and with optogenetic stimulation.**

Approximately 50 larvae at either St26 or St 29 chorionated larvae were transferred in 3cm uncoated petridishes. We incubated the plates (22 plates per larval stage) at 18°C in the dark. We checked each plate every 15 minutes and recorded the time point at which we identified the first settled larva. For statistical analysis we performed a Mann-Whitney test.

For the optogenetic experiments either St.26 larvae expressing Cii.Etr-1>CIBN-GFP-Caax; Cii.Etr-1>ARHGEF11(DHPH)-CRY2-mCherry or St.29 larvae expressing Cii.Etr-1>CIBN-mito-GFP;Cii.Etr-1>ARHGEF11(DHPH)-CRY2-mCherry. Around 50 transgenic larvae at either St.26 or St.29 were transferred to 3cm petri-dishes (coated briefly with 10mg/ml BSA) we repeated this experient in the presence of 10mM $NH_4Cl$, while illuminating with blue light (465–470 nm) using the LED Illumination Tool for Optogenetic Stimulation (LITOS) which is composed of an array of 232 × 64 LED matrix(*57*). We illuminated for 55 seconds, followed by 5 seconds of no light. We checked each plate every 15 minutes and recorded the time point at which we identified the first settled larva. For statistical analysis we performed a Mann-Whitney test.

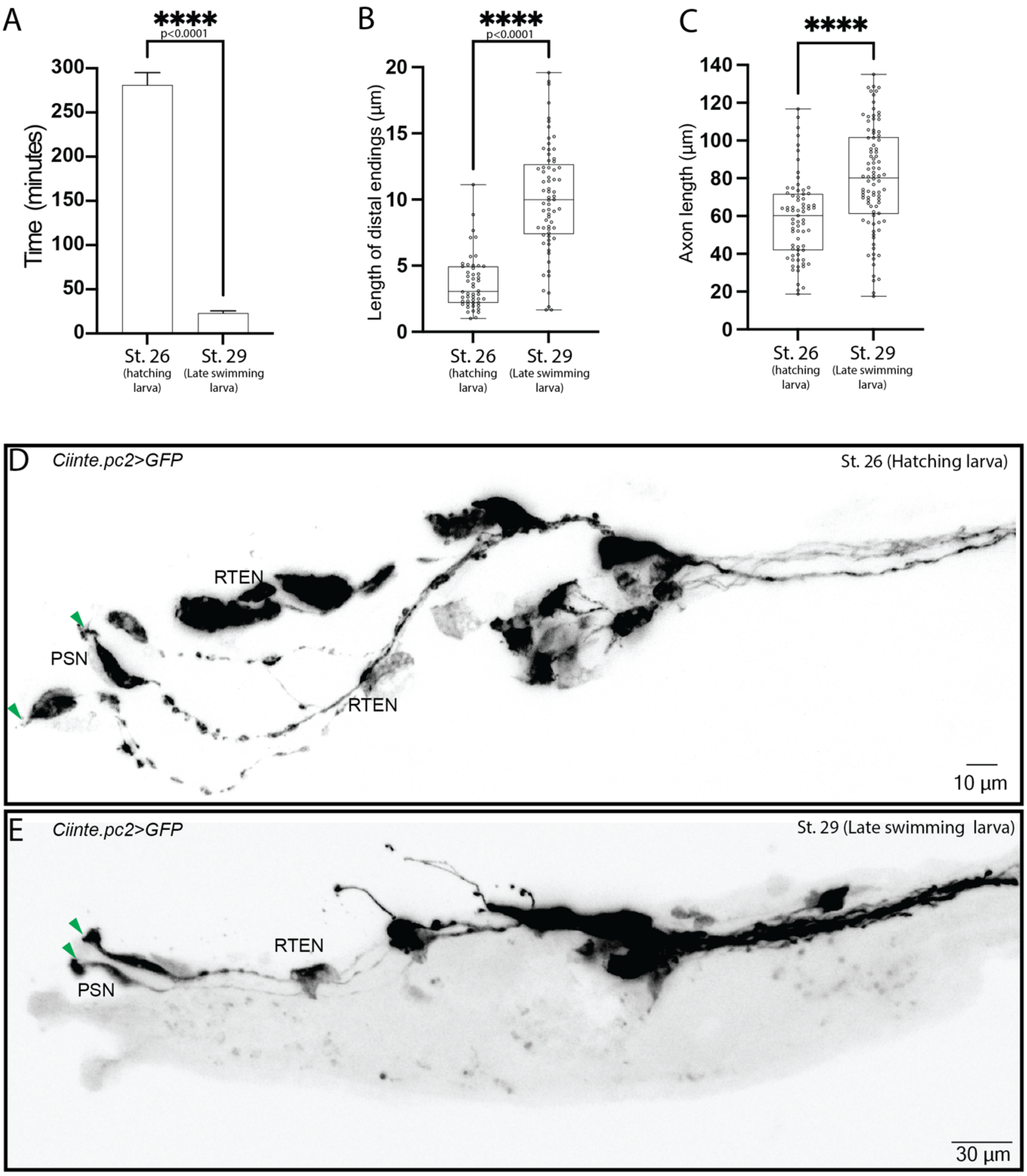


**Supplementary Figure S1: Late swimming larvae show enhanced settlement behavior and enhanced PSN cellular features compared to hatching larvae.**

(A) Quantification of settlement onset for freshly hatched larvae and late swimming larvae in the presence of 10mM $NH_4Cl$. 22 assay plates per stage, n≥50 per assay. For statistical analysis we performed a Mann-Whitney test. (B) Quantification of the distal sensory endings length in St. 26 hatching larvae (50 cells) and St. 29 late swimming larvae (69 cells). For statistical analysis we performed a Mann-Whitney test. (C) Quantification of the length of axons connecting the PSNs with the upstream RTENs neurons in St. 26 hatching larvae (71 cells) and St. 29 late swimming larvae (90 cells). For statistical analysis we performed a Mann-Whitney test. (D-E) Maximal projections of confocal stacks from Ciinte.pc2>GFP transgenic hatching larva (St. 26) (panel D) and late swimming larva (St. 29) (panel E). Scale bars are 10μm and 30μm respectively. Green arrowheads indicate the distal sensory endings of the PSNs.

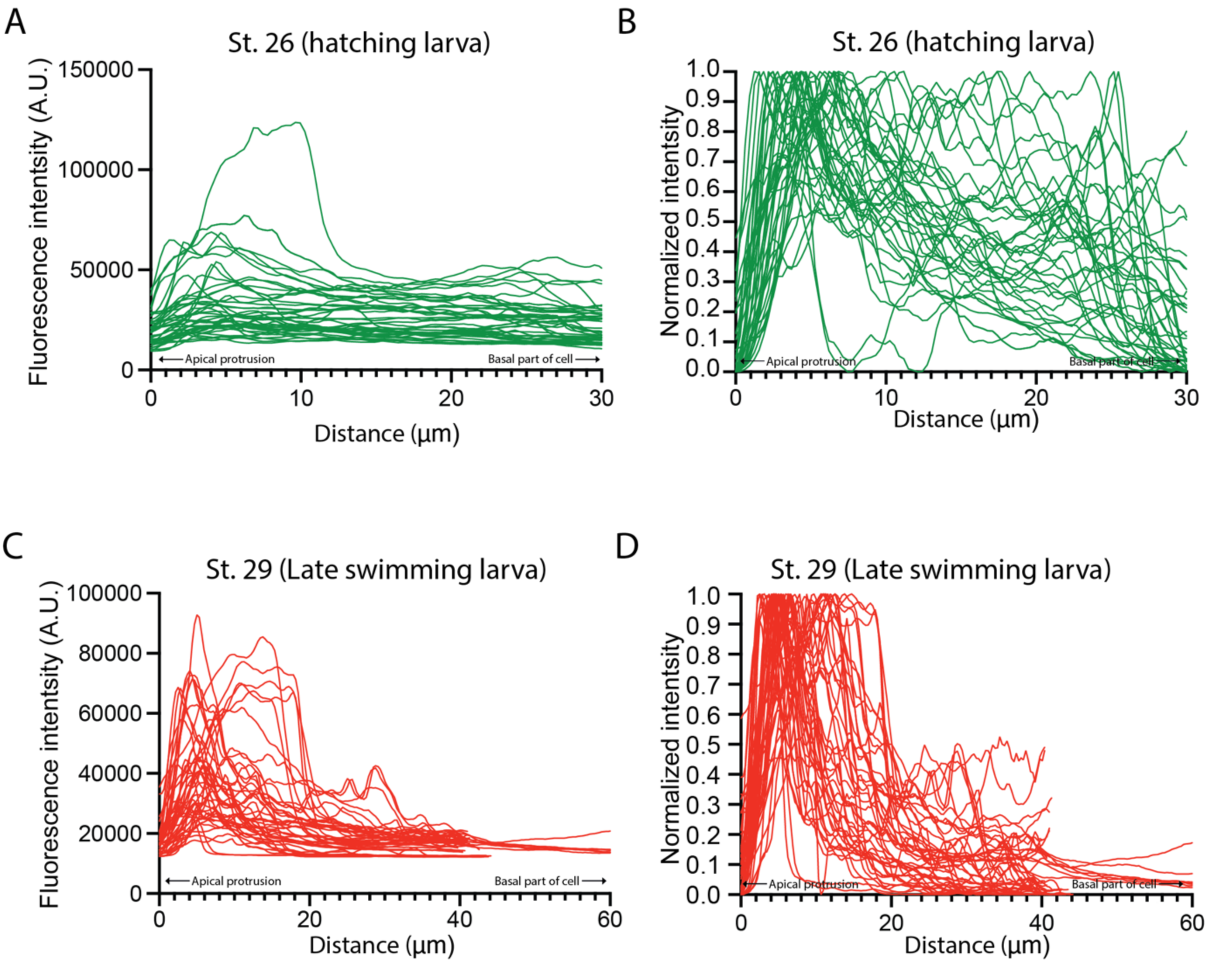


**Supplementary Figure S2: Utrophin localization in the**

(A-B) Raw intensity profiles (A) and corresponding normalized profiles (B) of individual papillae sensory cells expressing *mTurquoise::UtrCH* from St.26 hatching larvae (40 cells). (C-D) Raw intensity profiles and corresponding normalized profiles of individual papillae sensory cells expressing *mTurquoise::UtrCH* from St.29 late swimming larvae (56 cells). (A-D) underlie the average traces presented in panel Fig. 2H and 2I.

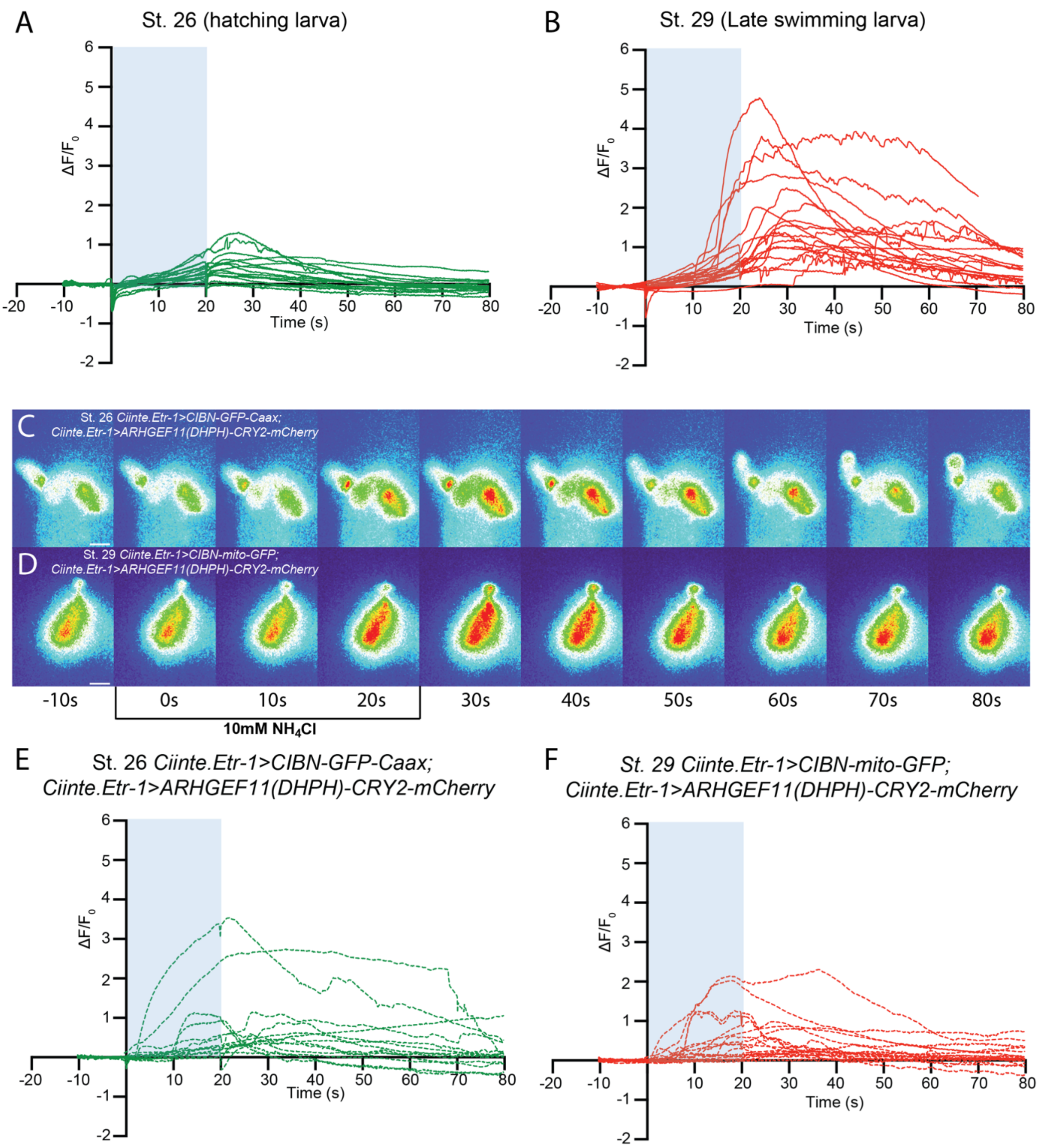


**Supplementary Figure S3: Chemosensory responses are enhanced in late swimming larvae and they can be modulated by the mechanical state of the sensory neurons.**

(A-B) Individual traces of Calcium responses to 10mM $NH_4Cl$ from the PSNs of (St.26) hatching larvae (panel A, 16 animals) and St. 29 late swimming larvae (panel B, 18 animals). Blue shaded region indicates stimulus delivery period. (C-D) Montage of calcium responses to 10mM $NH_4Cl$ exhibited by St. 26 *Ciinte.Etr-1>CIBN-GFP-Caax;Ciinte.Etr-1>ARHGEF11(DHPH)-CRY2-mCherry* (C) and St. 29 *Ciinte.Etr-1>CIBN-mito-GFP;Ciinte.Etr-1>ARHGEF11(DHPH)-CRY2-mCherry* (D). (E) (A-B) Individual traces of Calcium responses to 10mM $NH_4Cl$ from the PSNs of St. 26 *Ciinte.Etr-1>CIBN-GFP-Caax;Ciinte.Etr-1>ARHGEF11(DHPH)-CRY2-mCherry* larvae (panel E, 13 animals) and St. 29 *Ciinte.Etr-1>CIBN-mito-GFP;Ciinte.Etr-1>ARHGEF11(DHPH)-CRY2-mCherry* (panel F, 18 animals). Blue shaded region indicates stimulus delivery period. The individual traces presented in panels A, B, E and F underlie the average traces presented in panel Fig. 3C.

**Movie S1 Calcium response of St. 26 hatching larva PSNs responding to 10mM $NH_4Cl$**. Video of calcium activity in PSN neurons responding to 10mM $NH_4Cl$ from a St. 26 hatching transgenic larva expressing *Ciinte.pc2>GCaMP6s*.

**Movie S2 Calcium response of St. 29 late swimming larva PSNs responding to 10mM $NH_4Cl$**. Video of calcium activity in PSN neurons responding to 10mM $NH_4Cl$ from a St. 29 late swimming transgenic larva expressing *Ciinte.pc2>GCaMP6s*.

**Movie S3 Calcium response of St. 26 hatching larva PSNs responding to 10mM $NH_4Cl$. Expressing, combined with the activation of the optogenetic tool *CIBN-GFP-Caax; ARHGEF11(DHPH)-CRY2-mCherry.*** Video of calcium activity in PSN neurons responding to 10mM $NH_4Cl$ from a St. 26 hatching transgenic larva expressing *Ciinte.pc2>GCaMP6* and *Ciinte.Etr-1>CIBN-GFP-Caax;Ciinte.Etr-1>ARHGEF11(DHPH)-CRY2-mCherry.*

**Movie S4 Calcium response of St. 26 hatching larva PSNs responding to 10mM $NH_4Cl$. Expressing, combined with the activation of the optogenetic tool *CIBN-mito-GFP; ARHGEF11(DHPH)-CRY2-mCherry.*** Video of calcium activity in PSN neurons responding to 10mM $NH_4Cl$ from a St. 29 late swimming transgenic larva expressing *Ciinte.pc2>GCaMP6* and *Ciinte.Etr-1>CIBN-mito-GFP;Ciinte.Etr-1>ARHGEF11(DHPH)-CRY2-mCherry.*
